# The Interaction of *UBR4, LRP1*, and *OPHN1* in Refractory Epilepsy: *Drosophila* Model to Investigate the Oligogenic Effect on Epilepsy

**DOI:** 10.1101/2024.10.25.620163

**Authors:** Rui-Na Huang, Si-Yuan Luo, Tao Huang, Xiong-Sheng Li, Fan-Chao Zhou, Ling-Ying Li, Ze-Ru Chen, Sheng-Qiang Chen, Bin Tang, Jing-Da Qiao

## Abstract

Refractory epilepsy is an intractable neurological disorder that is currently difficult to achieve effective pharmacological control in clinical practice and can result in poor quality of life as well as increased mortality. Genetic factors are important causes of epilepsy, especially idiopathic epilepsy. In the clinical gene sequencing work, we identified one refractory epileptic patient who carried three epileptogenic candidate genes: *UBR4, LRP1*, and *OPHN1* variants. To clarify the epileptogenicity and interactions of *UBR4, LRP1*, and *OPHN1* variants, as well as explore the role of each mutant gene in eliciting epilepsy, we established single-knockdown, double-knockdown, and triple-knockdown *Drosophila* models by suppressing the gene expression of these three epileptogenic candidate genes. After conducting behavioral testing for epilepsy in the seven *Drosophila* knockdown models, regression equations illustrating the causal connection between genotype and phenotype were developed. The mutations of the three epileptogenic candidate genes: *UBR4, LRP1*, and *OPHN1*, were proved to be epileptogenic at the animal level both in seizure rates and logistic regression results. Moreover, significant negative interactions in *UBR4-OPHN1* KD and *LRP1-OPHN1* KD were found in the trigenic KD system as well as the *UBR4-OPHN1* and *LRP1-OPHN1* digenic KD system according to the logistic regression analysis result. However, despite the existence of negative interaction, three groups of digenic KD flies and one group of trigenic KD flies presented higher seizure rates than that of the corresponding monogenic KD flies. *LRP1-OPHN1* KD together with its negative interaction was regarded as the main causative factors for seizure in the *UBR4-LRP1-OPHN1* KD.

## Introduction

Epilepsy is a prevalent neurological condition characterized by a persistent propensity to spontaneous epileptic seizures, affecting almost 70 million individuals globally(Chen et al., 2017; Thijs et al., 2019). Over the past decade, the field of epilepsy genetics has advanced tremendously thanks to the advent of genomic technologies which expedited our comprehension of the genetic structure of epileptogenesis(Myers & Mefford, 2015; Perucca et al., 2020). Genetic factors are an important cause especially of idiopathic epilepsy, and de novo mutations have been demonstrated as a prominent genetic motive of serious seizures(Bhalla et al., 2011; Hildebrand et al., 2013). Despite a handful of inherited epilepsy instances being proved to be caused by monogenic mutation, the majority of cases are assumed to have a polygenic framework(Reid et al., 2009; Reid et al., 2010). With the widespread application of whole-exome sequencing technology in clinical diagnosis and cutting-edge genetic research, more candidate genes associated with epilepsy have been discovered(Takai et al., 2020). However, despite the availability of high-resolution and high-throughput sequencing methods, it is still difficult to elucidate the complex genetic pathogenic mechanism of refractory epilepsy. At present, most of the research on polygenic diseases focuses on bioinformatics analysis(Numis et al., 2022), and few studies on polygenic diseases are carried out at the whole animal level. Thus, elucidating the polygenic effects and interactions through direct experimentation is imperative for more precise diagnosis and targeted therapy especially given the current relative lack of polygenic research.

In most fields of research involving genetics and targeted therapies, the current researches on genetic mutations are often constructed at the cellular or tissue level. While differently in epilepsy, we verify the pathogenicity of mutations mainly at the animal level by observing epileptic behaviors. Drosophila melanogaster, which possesses elevated reproductive efficiency, outstanding versatility for genetic manipulation, and the nervous system that recapitulates a variety of cellular and network structures associated with human illnesses(Shulman, 2015), is ideal for confirming causal genes, illuminating their neurological impacts on seizures, and investigating novel biomarkers for drug-targeted treatment(Parker et al., 2011; Takai et al., 2020).

In this research, we analyzed the causative genes of 436 patients with idiopathic epilepsy by trio-based whole-exome sequencing (WES) and identified one child patient with serious manifestations carrying three potentially epileptogenic genetic mutations (UBR4: missense mutation; LRP1: missense mutation; OPHN1: shearing mutation). Utilizing the UAS-RNAi/ Gal4 system and double balancer tool, we constructed genetic knockdown (KD) models in Drosophila which were comprised of Poe (the ortholog of UBR4), LRP1 and Graf (the ortholog of OPHN1) single KD flies, three double KD flies as well as one triple KD flies. To clarify the epileptogenicity and interactions of the three genes, we conducted Bang-Sensitive (BS) tests to obtain the seizure rates of the 7 types of flies and the controls separately. Then, we constructed logistic regression equations based on these statistics data to investigate the relationship between genotypes and phenotypes and lastly elucidate the main causative factor(s) by mathematical quantification for carrying out the targeted pharmacological tests.

## Materials and methods

### 1. Whole-exome sequencing and Clinical-Genetic-Database synthetical analysis

Idiopathic epilepsy cases were obtained from the Epilepsy Center of the Second Affiliated Hospital of Guangzhou Medical University. Meanwhile, detailed clinical data including family history, age at seizure, seizure type and frequency, seizure course, neurological examination results such as EEG, and antiepileptic treatment history were collected. Peripheral blood samples were obtained from patients and their parents or other family members as available to determine the source of the identified genetic variants. DNA fragments from the whole exon region of the human genome were captured using NanoWES probes. Whole exon high throughput sequencing was performed using the Novaseq 6000 sequencing platform. Positive loci were verified by Sanger sequencing using the ABI 3730 sequencing platform. Genetic phenotype information was obtained from the OMIM database. SIFT, Polyphen2, MutationTaster, MutationAssessor, FATHMM, PROVEAN, MetaSVM, MetaLR, and LRT databases were used to predict the deleteriousness of genetic variants. The protein fragment structure and DeltaDeltaG value of the missense mutations were predicted using Alphafold3 and I-mutant 2.0(Abramson et al., 2024; Capriotti et al., 2005). Figures of the structure model are processed with PyMol 3.0.5 software.

### 2. Knockdown Drosophila models generation

The tub-Gal4 line was hybridized with UBR4-RNAi, LRP1-RNAi, and OPHN1-RNAi lines separately to obtain UBR4, LRP1, and OPHN1 single knockdown flies (tub-Gal4; UBR4-RNAi, tub-Gal4; LRP1-RNAi, and tub-Gal4; OPHN1-RNAi)(Liu et al., 2023). In combination with a double balancer line (Cyo; Gla/TM3; Ser), UBR4-LRP1, UBR4-OPHN1 and LRP1-OPHN1 double knockdown flies (tub-Gal4; UBR4-RNAi/ LRP1-RNAi, tub-Gal4/ OPHN1-RNAi; UBR4-RNAi,, and tub-Gal4/ OPHN1-RNAi; LRP1-RNA;), as well as UBR4-LRP1-OPHN1 triple knockdown flies (tub-Gal4/ OPHN1-RNAi; UBR4-RNAi/ LRP1-RNAi;) can be established.

### 3. RT-qPCR

To ascertain the expression levels of UBR4, LRP1, and OPHN1 in knockdown models, RT-qPCR was carried out as described below. In short, eight post-eclosion flies (Canton-S (WT), tub-Gal4; UBR4-RNAi, tub-Gal4; LRP1-RNAi, tub-Gal4/ OPHN1-RNAi, tub-Gal4; UBR4-RNAi/ LRP1-RNAi, tub-Gal4/ OPHN1-RNAi; UBR4-RNAi;, tub-Gal4/ OPHN1-RNAi; LRP1-RNAi;, and tub-Gal4/ OPHN1-RNAi; UBR4-RNAi/ LRP1-RNAi) were frozen at −80°C shortly after being isolated. RNA was decontaminated utilizing HiPure Universal RNA Mini Kit (Magen Biotechnology, Guangzhou), cDNA was compounded utilizing HiScript III RT SuperMix (Nanjing Vazyme Biotech Co., Ltd., Nanjing) and qPCR was executed utilizing Taq Pro Universal SYBR qPCR Master Mix (Nanjing Vazyme Biotech Co., Ltd., Nanjing). For the synthesis of cDNA, random decamers were employed as primers. Afterward, transcripts in each genetic background were identified by amplifying the cDNA of UBR4, LRP1, and OPHN1. The following primers were used:

UBR4/Poe-F1: TCCGATGATGATGAAGGTTCCAATG

UBR4/Poe-R1: CTACTCCACTGCCTCCACTCTC

LRP1-F1: TGCGGCGATGGTAGTGATGAG

LRP1-R1: CAGAGGAACTTGCGAGGAATGC

OPHN1/Graf-F1: TGAGACCAGCACACGACGATAC

OPHN1/Graf-R1: CGCCGTTGTTAGCACCTATAATACC

### 4. Seizure behavior test

The seven types of knockdown flies, together with the UBR4-RNAi (control), LRP1-RNAi (control), and OPHN1-RNAi (control) flies were all submitted to a seizure behavior test. The bang-sensitive (BS) test was performed on flies 3–5 days after eclosion and seizure-like behavior was assessed. A day before testing, the flies were transferred in fresh and clean feeding vials after receiving CO2 anesthesia. About three to six flies were contained per vial and mechanically irritated at maximum speed for 20s using a vortex mixer (VWR) (Liu et al., 2023). Seizure behaviors in flies were recorded in terms of both percentage and continuance. Each genotype was assessed through at least seven trials which consisted of five to seven vials of flies.

### 5. Logistic regression (LR) models

To ascertain the effect of each gene’s KD and their interactions on seizures in the polygenic system (Stoltzfus, 2011), we established four LR models for quantitative analyses employing SPSS 25.0 software. According to the binary dependent variable applied in the LR model, “seizure” was coded as 1, and “no seizure” was coded as 0.

Firstly, in the trigenic model based on the patient genotype, we assumed three covariates X_1_, X_2_, and X_3_ respectively (where KD = 1 signifies gene knockdown and KD = 0 signifies no knockdown, Table 3) in the trigenic model to represent UBR4 KD, LRP1 KD, and OPHN1 KD based on the odds of seizure outcome. The equation for this model was as followed:

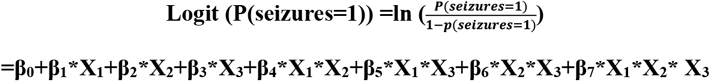

**Table 1.** Whole Exome Sequencing results (Identification of *UBR4, LRP1*, and *OPHN1* variants)

| Mutant Gene | Mutation Location Transcript (hg38) | Heterozygosity | Variation Type | Other Sample Results |  |
| --- | --- | --- | --- | --- | --- |
|  |  |  |  | Father | Mother |
| *UBR4* | chr1: 19160975<br>NM_020765.3 | heterozygous | missense variation | heterozygous | negative |
| *UBR4* | chr1: 19150673<br>NM_020765.3 | heterozygous | missense variation | negative | heterozygous |
| *LRP1* | chr12: 57193256<br>NM_002332.3 | heterozygous | missense variation | heterozygous | negative |
| *LRP1* | chr12: 57201042<br>NM_002332.3 | heterozygous | missense variation | negative | heterozygous |
| *OPHN1* | chrX: 68064181<br>NM_002547.3 | heterozygous | shearing variation | negative | negative |

**Table 2.** Comprehensive analysis outcomes based on genetics and database information.

| Mutant Gene | Biological Hazard <sup>1</sup> | Population Frequency (MAF) <sup>5</sup> |  |  |  | OMIM |  | ACMG Pathogenicity <sup>2</sup> | Clinical Consistency Assessment |
| --- | --- | --- | --- | --- | --- | --- | --- | --- | --- |
|  |  | KG | gnomAD | gnomAD-EAS | gnomAD-Hom | Disease Name <sup>3</sup> | Hereditary Mode <sup>4</sup> |  |  |
| *UBR4* | 3 | - | 0.00000<br>6575 | 0.0001928 | 0 | - | - | US | possible pathogenicity |
| *UBR4* | 3 | - | - | - | - |  |  | US |  |
| *LRP1* | 6 | 0.002<br>415 | 0.00308<br>7000 | 0.0021250 | 2 | KPA | AR | US | possible pathogenicity |
| *LRP1* | 6 | - | 0.00001<br>3150 | - | 0 |  |  | US |  |
| *OPHN1* | - | - | - | - | - | Mental retardation, X-linked, with cerebellar hypoplasia and distinctive facial appearance | XLR | LP | possible pathogenicity |

1. The number indicates the number of databases (SIFT, Polyphen2, MutationTaster, MutationAssessor, FATHMM, PROVEAN, MetaSVM, MetaLR, LRT, etc.) in which the deleteriousness of a gene variant was assessed.
2. The determination of pathogenicity of genetic variants was based on the criteria for interpretation of sequence variants published by the American College of Medical Genetics and Genomics (ACMG), which were categorized as follows: Pathogenic; LP (Likely pathogenic); US (Uncertain significance); Likely benign; Benign.
3. AFND (Acromelic frontonasal dysostosis); NEDMAGA (Neurodevelopmental disorder with movement abnormalities, abnormal gait, and autistic features); KPA (Keratosi pilaris atrophicans).
4. AD, autosomal dominant; AR, autosomal recessive; XL, X chromosome; XLD, X chromosome dominant; XLR, X chromosome recessive.
5. KG (1000 Genomes Project); gnomAD (The Genome Aggregation Database); gnomAD-EAS (EAS for East Asians); gnomAD- Hom (number of homozygotes)

**Table 3.**
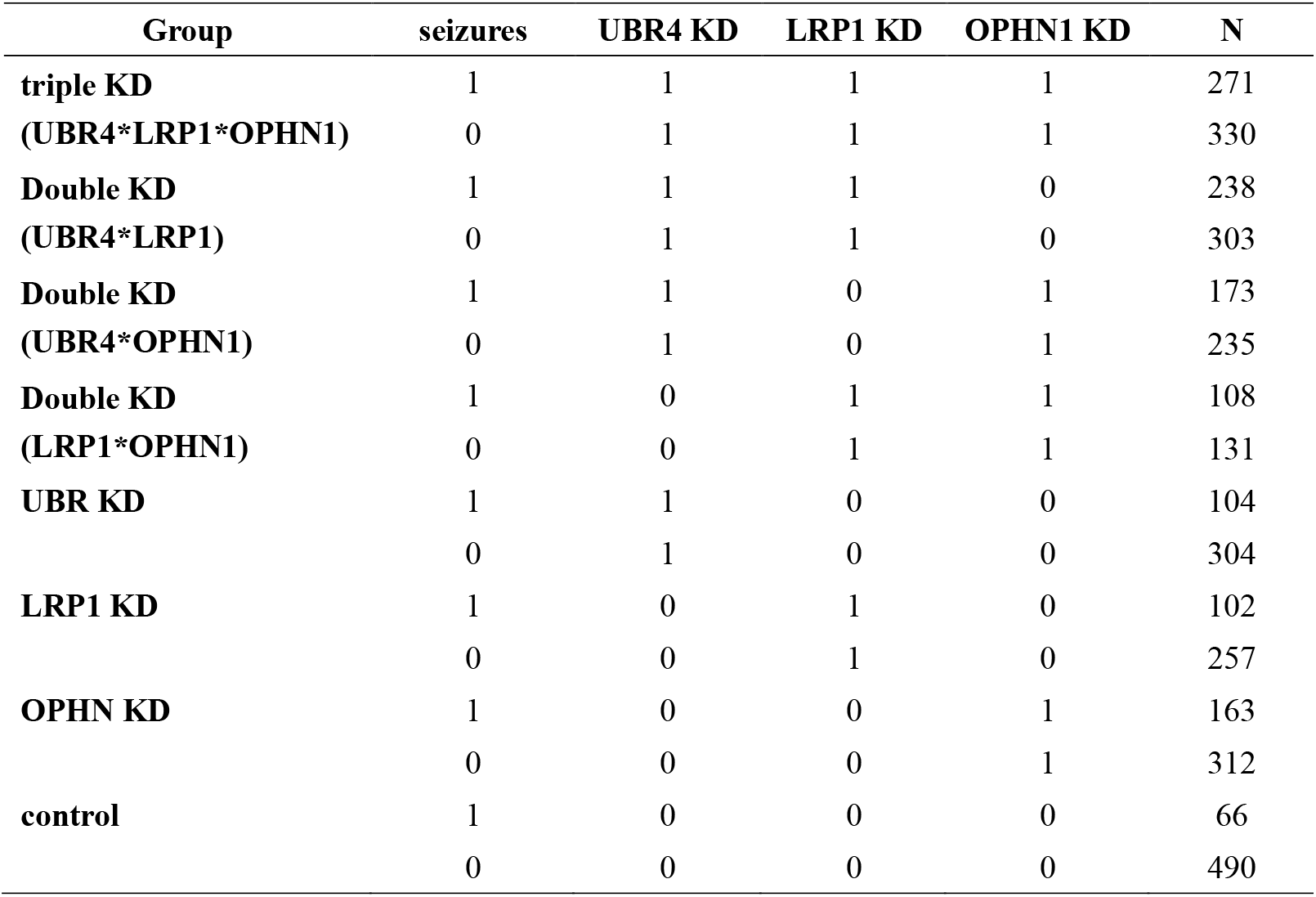
Total seizures and the KD of three genes (*UBR4*&*LRP1*&*OPHN1*)

| Group | seizures | UBR4 KD | LRP1 KD | OPHN1 KD | N |
| --- | --- | --- | --- | --- | --- |
| **triple KD** | 1 | 1 | 1 | 1 | 271 |
| **(UBR4\*LRP1\*OPHN1)** | 0 | 1 | 1 | 1 | 330 |
| **Double KD** | 1 | 1 | 1 | 0 | 238 |
| **(UBR4\*LRP1)** | 0 | 1 | 1 | 0 | 303 |
| **Double KD** | 1 | 1 | 0 | 1 | 173 |
| **(UBR4\*OPHN1)** | 0 | 1 | 0 | 1 | 235 |
| **Double KD** | 1 | 0 | 1 | 1 | 108 |
| **(LRP1\*OPHN1)** | 0 | 0 | 1 | 1 | 131 |
| **UBR KD** | 1 | 1 | 0 | 0 | 104 |
|  | 0 | 1 | 0 | 0 | 304 |
| **LRP1 KD** | 1 | 0 | 1 | 0 | 102 |
|  | 0 | 0 | 1 | 0 | 257 |
| **OPHN KD** | 1 | 0 | 0 | 1 | 163 |
|  | 0 | 0 | 0 | 1 | 312 |
| **control** | 1 | 0 | 0 | 0 | 66 |
|  | 0 | 0 | 0 | 0 | 490 |

Here, β_0_ is the intercept; β_1_, β_2_, β_3_, β_4_, β_5_, β_6_, β_7_ are the regression coefficients for the independent variables X_1_, X_2_, X_3_, X_1_*X_2_, X_1_*X_3_, X_2_*X_3_ and X_1_*X_2_* X_3_

Moreover, we assumed two covariates X_1_ and X_2_, respectively (where KD= 1 signifies gene knockdown and KD= 0 signifies no knockdown) to represent the corresponding monogenic KD in the three digenic models (Supplementary table 1, table 4, table 7). The following was the equation for each of these three models:

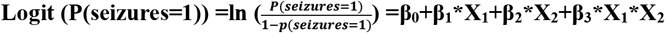

**Table 4.** LR outcomes (interaction among *UBR4, LRP1* and *OPHN1*)

|  | B | S.E | Wald | OR | 95% CI for |  | P value |
| --- | --- | --- | --- | --- | --- | --- | --- |
|  |  |  |  |  | Odds |  |  |
|  |  |  |  |  | Lower | Upper |  |
| UBR4 KD | 0.932 | 0.173 | 28.868 | 2.540 | 1.808 | 3.568 | 0.000*** |
| LRP1 KD | 1.081 | 0.176 | 37.808 | 2.947 | 2.088 | 4.158 | 0.000*** |
| OPHN1 KD | 1.355 | 0.163 | 69.250 | 3.879 | 2.819 | 5.337 | 0.000*** |
| UBR4*LRP1 | -0.249 | 0.226 | 1.213 | 0.779 | 0.500 | 1.215 | 0.271ns |
| UBR4*OPHN1 | -0.589 | 0.222 | 7.016 | 0.555 | 0.359 | 0.858 | 0.008** |
| LRP1*OPHN1 | -0.624 | 0.239 | 6.827 | 0.536 | 0.335 | 0.856 | 0.009** |
| UBR4*LRP1*OPHN1 | -0.097 | 0.307 | 0.101 | 0.907 | 0.497 | 1.656 | 0.751ns |
| Constant | -2.005 | 0.131 | 233.768 | 0.135 |  |  | 0.000*** |

Here, β_0_ is the intercept; β_1_, β_2_, and β_3_ are the regression coefficients for the independent variables X_1_, X_2_, and X_1_*X_2_.

### 6. Statistical analysis

All quantitative data were presented as Mean ± Standard Error of the Mean (SEM). Two independent or paired samples were compared employing the Student’s t-test. One-way ANOVA was utilized to analyze multiple samples, and Tukey’s post hoc test was used to evaluate differences between the two groups. Seizure behavior data were also analyzed by Chi-square test. Statistical analyses were conducted with GraphPad Prism 9.5 and SPSS 25.0. The cut-off value for statistical significance was 0.05.

## Results

### 1. Identification of UBR4, LRP1 and OPHN1 variants

In an idiopathic refractory epilepsy case, heterozygous variants in three genes, UBR4, LRP1, and OPHN1, were simultaneously identified. The patient is a female with no familial history of epilepsy, who started at the age of 4 years with complex partial seizures (CPS) type characterized by partial seizures, and relapsed after surgical treatment with microdissection of epileptic foci at the age of 5 years, accompanied by simple partial seizures (SPS). Her seizures possessed a maximum frequency of up to 20 per month, each lasting 30 seconds to 1 minute, and EEG showed abnormal EEG slowing. In the early stages of the disease, seizure-free treatment for 3 months could be achieved after Levetiracetam monotherapy, however, after postoperative relapse, effective seizure control could not be achieved with Lacosamide alone and even with a combination of six antiepileptic drugs: LEV (Levetiracetam)+ LTG (Lamotrigine)+ VPA (Valproic Acid)+ OXC (Oxcarbazepine)+ LCM (Lacosamide)+ CNZ (Clonazepam).

Among these three genes, UBR4 (chr1: 19160975/NM_020765.3 and chr1: 19150673/ NM_020765.3) includes two missense variants, LRP1 (chr12: 57193256/NM_002332.3 and chr12: 57201042/ NM_002332.3) includes two missense variants, and OPHN1 (chrX: 68064181/NM_002547.3) includes one shear variant. Notably, the OPHN1 gene shear variant is a de novo variant of nonparental origin and was not listed in any public databases (including 1000 Genomes Project and gnomAD). The variants of UBR4 gene were predicted to be biohazardous in three of the SIFT, Polyphen2, MutationTaster, MutationAssessor, FATHMM, PROVEAN, MetaSVM, MetaLR, LRT databases, etc. and LRP1 was predicted to be biohazardous in six databases. Based on the criteria for interpretation of sequence variants published by the American College of Medical Genetics and Genomics (ACMG), the pathogenicity of UBR4, LRP1, and OPHN1 genetic variants was of uncertain significance, but were all considered potentially pathogenic in the clinical concordance assessment. (Tables 1, 2) To investigate the functional impact of missense mutations in UBR4 and LRP1, we predicted the changes in the local structures of the wild-type and mutant proteins. The results revealed that the hydrogen bonding structure between amino acids did not change before and after the mutation, however, the DeltaDeltaG value of the four missense variants were all negative, suggesting the decrease in the stability of the protein structure (Figure 1). Accordingly, we hypothesized that these four missense mutations are LOF (Loss-Of-Function) mutations, which are applicable to the knockdown Drosophila model for simulation.

**Figure 1.**
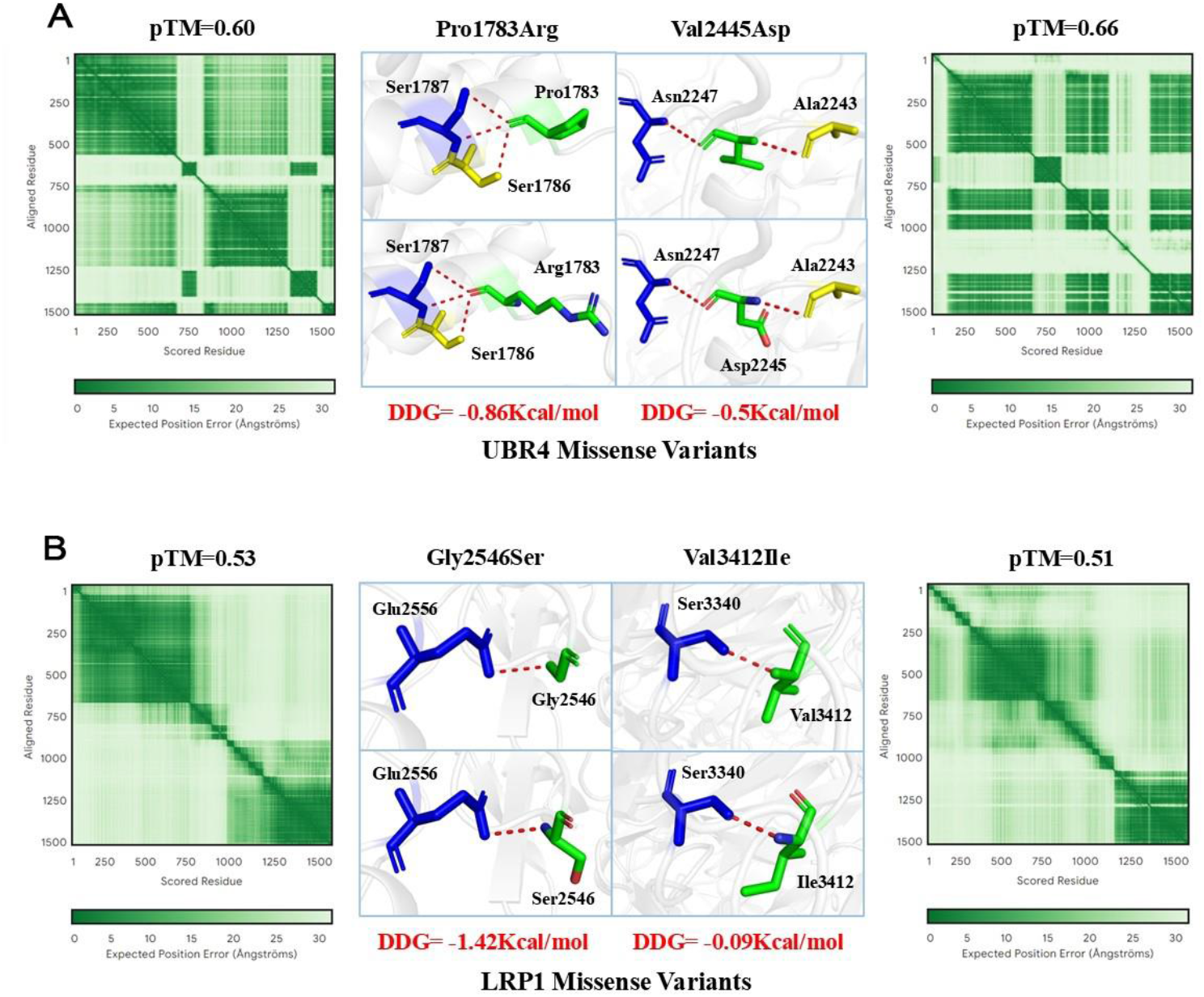
Protein structure prediction for missense mutations in UBR4 and LRP1. (A) Localized protein structure prediction at the UBR4 missense mutation site. (B) Localized protein structure prediction at the LRP1 missense mutation site.

### 2. RT-qPCR

As the results presented in Figure 2, UBR4, LRP1, and OPHN1 were KD successfully in seven drosophila strains. The KD efficiency of tub-Gal4>UBR4-RNAi, tub-Gal4>LRP1-RNAi and tub-Gal4>OPHN1-RNAi was 78.30%, 69.40% and 75.90%, separately [0.2170±0.0345% (n=3) versus 1.000±0.1470% (n=3), **P=0.007; 0.3060±0.0454% (n=4) versus 1.0000±0.1870% (n=3), **P=0.009; 0.2410±0.0828% (n=2) versus 1.0000±0.0260% (n=3), **P=0.002. Figure 2 A, B, C]. Moreover, the relative mRNA expression level of the genes was all significantly lower than the controls in the digenic and trigenic KD flies, indicating that drosophila models with polygenic KD were successfully constructed.

**Figure 2.**
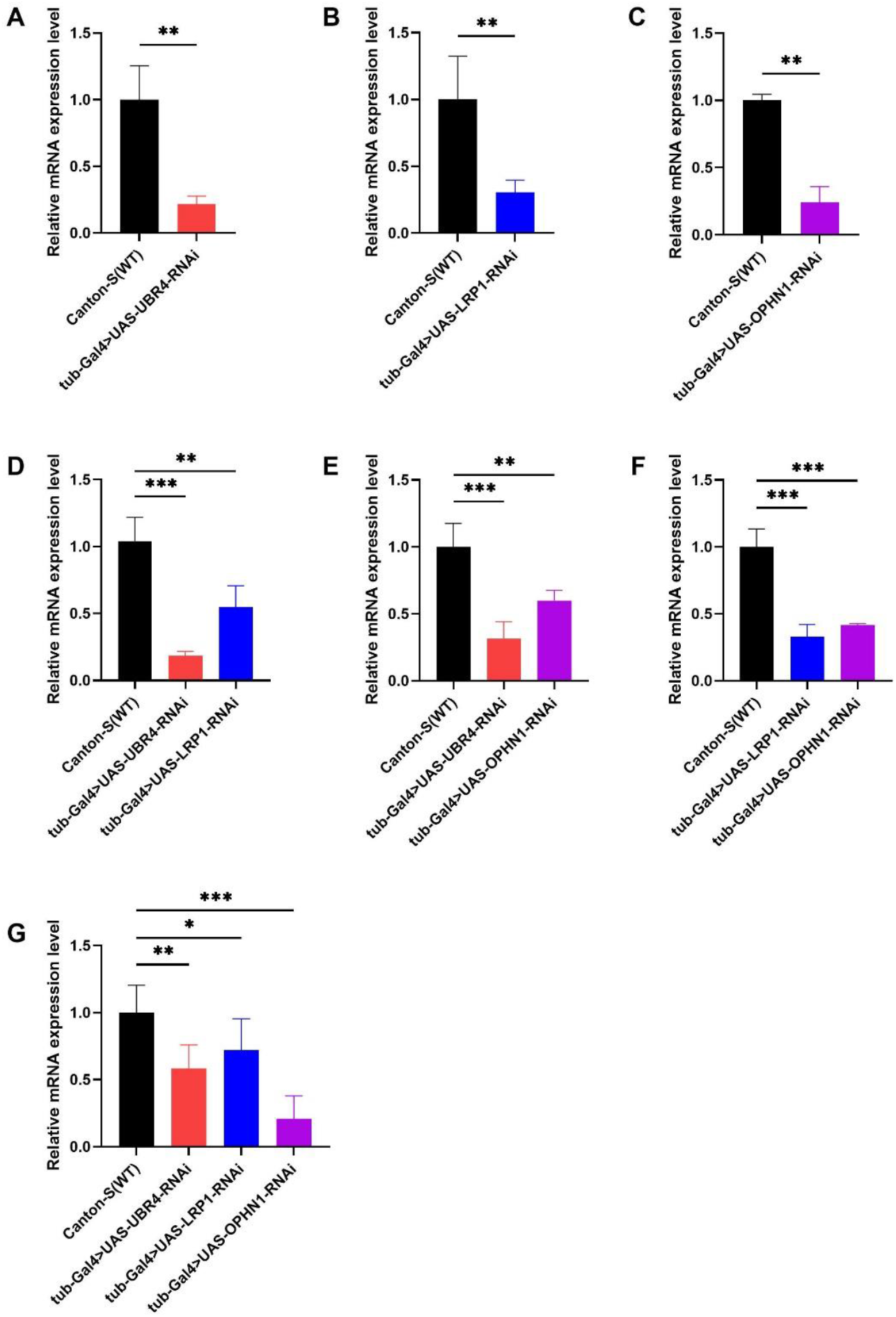
KD efficiency of seven fly strains. (A) Relative expression of mRNA in *UBR4* KD flies and wild-type flies. (B) Relative expression of mRNA in *LRP1* KD flies and wild-type flies. (C) Relative expression of mRNA in *OPHN1* KD flies and wild-type flies. (D) Relative expression of mRNA in *UBR4, LRP1* double KD flies, and wild-type flies. (E) Relative expression of mRNA in *UBR4, OPHN1* double KD flies, and wild-type flies. (F) Relative expression of mRNA in *LRP1, OPHN1* double KD flies, and wild-type flies. (G) Relative expression of mRNA in *UBR4, LRP1, OPHN1* triple KD flies, and wild-type flies.

### 3. Seizure behavior of knockdown flies

In addition to classical seizure behavior, we observed a judder seizure behavior characterized by high-frequency flapping of the wings and a hyperdynamic seizure behavior that combines both judder and classical seizures, when performing the BS test on the three double KD and the triple KD drosophila models. In our recent study, flies of the well-known gene CAC KD, the candidate gene CELSR2 KD, and the double KD of these two genes also presented similar behavior. We hypothesize that these two more intense behaviors may be associated with genetically greater epileptogenicity.

#### 3.1. Monogenic KDs of UBR4, LRP1 and OPHN1 were revealed to be epileptogenic

Flies crossing strategy in seizure behavior test of the monogenic KDs was shown in Figure 3 A. The seizure rates of UBR4 KD flies, LRP1 KD flies and OPHN1 KD flies were discovered to be significantly higher than those of the controls [tub-Gal4>UBR4-RNAi, 25.490% (n=408) versus UBR4-RNAi, 12.705% (n=244), ***P<0.001; tub-Gal4>LRP1-RNAi, 28.412% (n=359) versus LRP1-RNAi, 10.612% (n=245), ***P<0.001; tub-Gal4>OPHN1-RNAi, 34.316% (n=475) versus OPHN1-RNAi, 13.433% (n=67), ***P<0.001. Figure 3 B, C, D]. Additionally, the recovery time of LRP1 KD and OPHN1 KD flies was longer than that of the controls. Most tub-Gal4>LRP1-RNAi flies recovered within 1-5s, whereas LRP1-RNAi flies recovered within 1-2s (Figure 3 C). The majority of tub-Gal4>OPHN1-RNAi flies recovered in more than 10s, while OPHN1-RNAi flies recovered within 1-4s (Figure 3 D). It pointed out that the monogenic knockdown variants of these three candidate genes were all associated with epilepsy.

**Figure 3.**
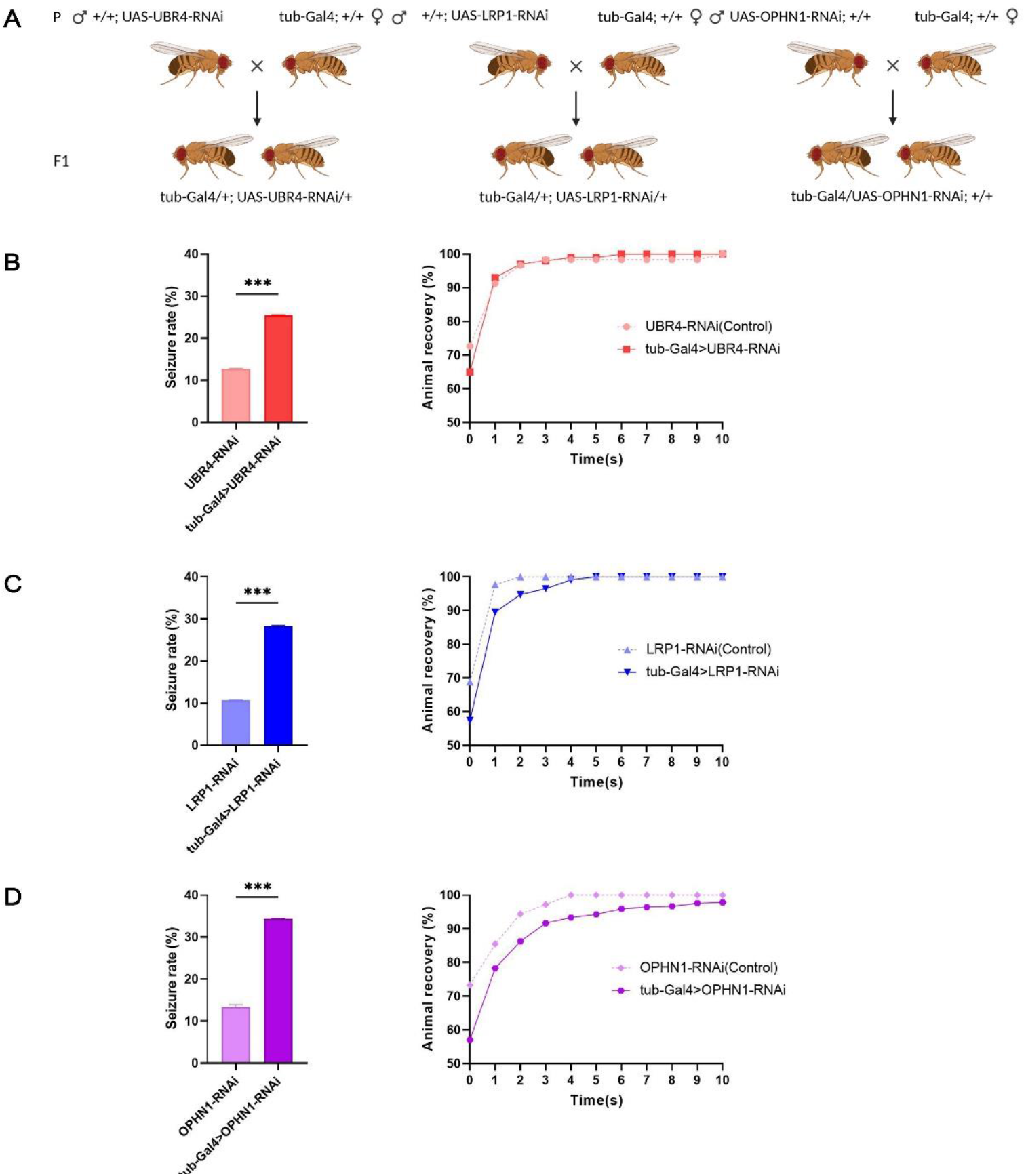
Pathogenicity of *UBR4, LRP1* and *OPHN1* variants. (A) Experimental procedure of flies crossing. (B)Seizures rates and recovery time of *UBR4* KD flies together with the controls. (C) Seizure rates and recovery time of *LRP1* KD flies together with the controls. (D) Seizure rates and recovery time of *OPHN1* KD flies together with the controls. Error bars represent 95% Confidence Interval (CI) of seizure rates.

### 3.2. LRP1-OPHN1 KD was major epileptogenic factor in the trigenic KD system

Crossing strategies of multigenetic KD flies in seizure behavior test were shown in Figure 4 A. As for the UBR4-LRP1-OPHN1 triple KD flies, 45.092% of them showed seizure behavior, which was significantly higher than that of the single KD flies [tub-Gal4>UBR4-RNAi; LRP1-RNAi, OPHN1-RNAi, 45.092% (n=601) versus tub-Gal4>UBR4-RNAi, 25.490% (n=408), ***P<0.001; versus tub-Gal4>LRP1-RNAi, 28.412% (n=359), ***P<0.001; versus tub-Gal4>OPHN1-RNAi, 34.316% (n=475), ***P<0.001. Figure 4 B]. Moreover, the seizure rate of OPHN1 KD flies was significantly higher than UBR4 KD flies [tub-Gal4>OPHN1-RNAi, 34.316% (n=475) versus tub-Gal4>UBR4-RNAi, 25.490% (n=408), **P=0.004. Figure 4 B]. The recovery time of UBR4-LRP1-OPHN1 KD flies was longer than that of UBR4 KD and LRP1 KD flies. The majority of tub-Gal4>UBR4-RNAi; LRP1-RNAi; OPHN1-RNAi together with tub-Gal4>OPHN1-RNAi flies recovered in more than 10s, while tub-Gal4>UBR4-RNAi flies recovered within 1-6s and tub-Gal4>LRP1-RNAi flies recovered within 1-5s (Figure 4 B). It suggested that trigenic KD were more promotive on seizures compared with monogenic KDs and OPHN1 KD flies showed more intense epileptic behavior. However, there was no apparent difference in seizure rate and recovery time when comparing trigenic KD flies to digenic KD flies, except for the slower recovery rate of tub-Gal4>LRP1-RNAi; OPHN1-RNAi flies (Figure 4 C).

**Figure 4.**
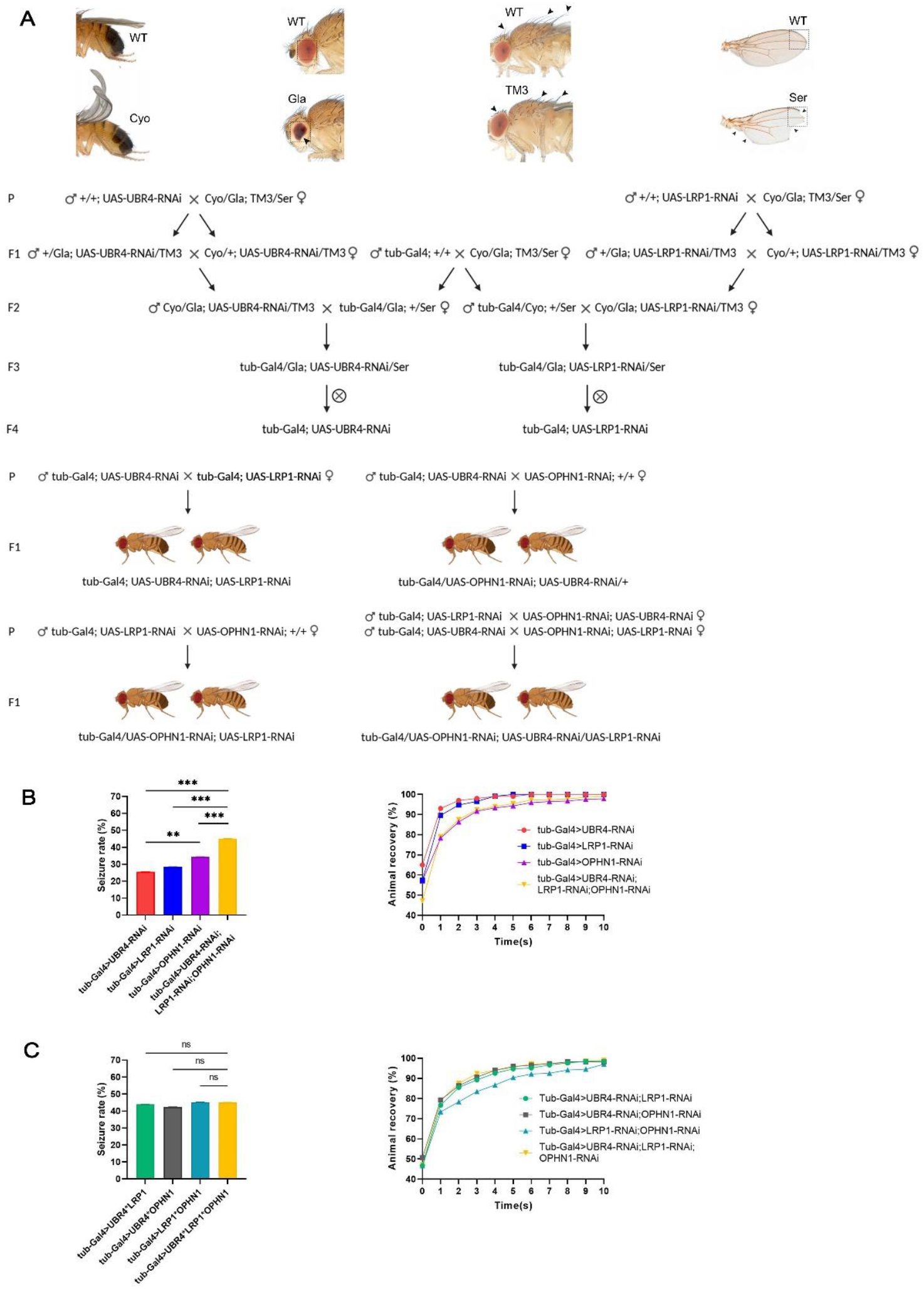
Seizure behavior of trigenic KD flies compared with monogenic KD and digenic KD flies. (A) Experimental procedure of flies crossing. (B)Seizure rates and recovery time of *UBR4* KD, *LRP1* KD, *OPHN1* KD and *UBR4-LRP1-OPHN1* KD flies. (B) Seizure rates and recovery time of *UBR4-LRP1* KD, *UBR4-OPHN1* KD, *LRP1-OPHN1* KD and *UBR4-LRP1-OPHN1* KD flies. Error bars represent 95% Confidence Interval (CI) of seizure rates.

According to the LR results of trigenic KD system in Table 4, the B value and OR value of UBR4 KD, LRP1 KD, and OPHN1 KD were separately 0.932, 2.540 (***P<0.001), 1.081, 2.947 (***P<0.001), and 1.355, 3.879 (***P<0.001), illustrating that every monogenic KD was a seizure risk factor in the trigenic KD system (Figure 5A). Besides, we found that UBR4*OPHN1 KD and LRP1*OPHN1 KD exhibited negative B values as −0.589 (**P=0.008) and −0.624 (**P=0.009), suggesting the significant negative interactions between UBR4 and OPHN1, as well as LRP1 and OPHN1, which signifies the protective effects against seizure beyond the multiplicative effect of the two genes. However, the ROC curve based on the LR results showed that the combination of the three genes predicted seizures with higher accuracy (Figure 5B). The LR equation with a bit less significance for this model was as follows:

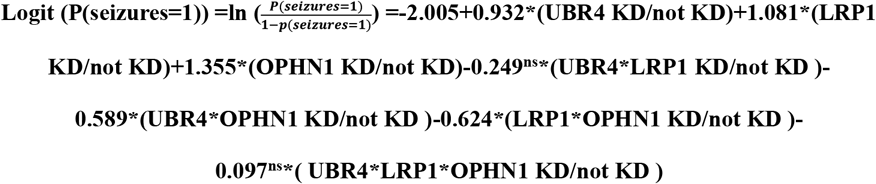

**Figure 5.**
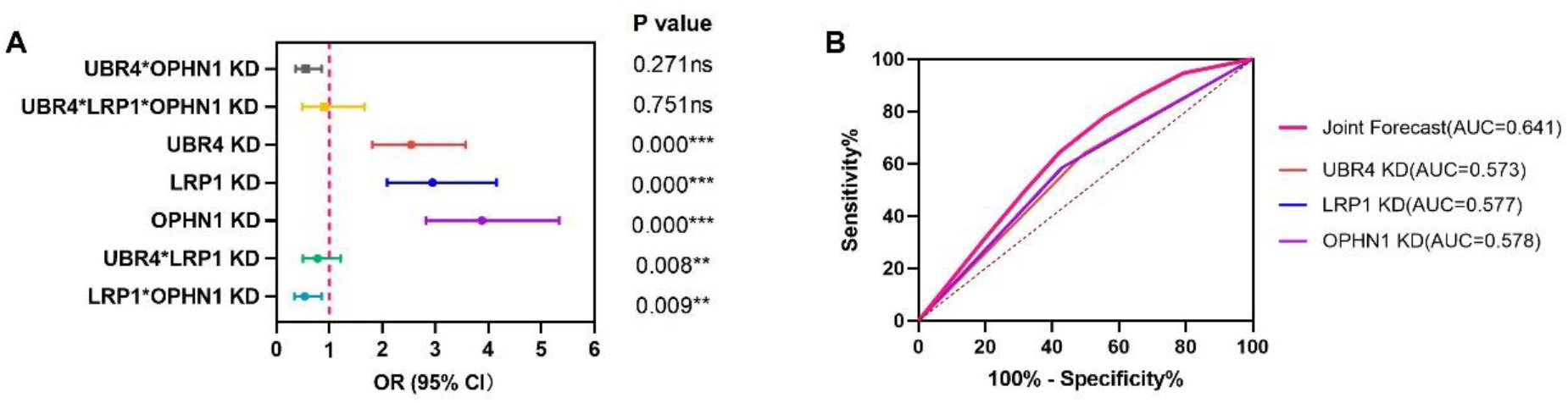
LR analysis of the trigenic KD system. (A)Forest plot of Odds Ratio(OR) for the KDs. (B)Receiver Operating Characteristic (ROC) Curve of UBR4 KD, LRP1 KD, OPHN1 KD, and the Joint Forecast by Logistic regression.

To further determine the influence of genes in the trigenic KD system, the variates of the LR equation were analyzed in more detail. Considering the insignificant relation to the monogenic KDs on seizures in Table 5, we additionally analyzed the effects of the digenic KDs together with their corresponding significant or insignificant interactions. As displayed in Table 6, when another gene was KD, the B value of the digenic KDs was noticeably smaller than that of not KD [OPHN1KD 0.226 versus OPHN1 not KD 1.110; LRP1 KD 0.236 versus OPHN1 not KD 0.894; UBR4 KD 0.295 versus UBR4 not KD 0.970]. While the positive value still indicated the existent but reduced epileptogenic effects when three genes were all knockdown. Based on the larger B value of LRP1 KD, OPHN1 KD, and LRP1-OPHN1 KD, the digenic KD of LRP1 and OPHN1 together with its significant negative interaction was regarded as the main factor on seizures in the trigenic KD system.

**Table 5.** LR outcomes (seizures in relation to the monogenic KDs)

|  |  | B | S.E | Wald | OR | 95% CI for Odds |  | P value |
| --- | --- | --- | --- | --- | --- | --- | --- | --- |
|  |  |  |  |  |  | Lower | Upper |  |
| LRP1-OPHN1<br>was KD | UBR4 KD | -0.004 | 0.154 | 0.001 | 0.996 | 0.737 | 1.346 | 0.980ns |
|  | Constant | -0.193 | 0.130 | 2.207 | 0.824 |  |  | 0.137ns |
| UBR4-OPHN1<br>was KD | LRP1 KD | 0.109 | 0.129 | 0.713 | 1.116 | 0.866 | 1.438 | 0.398ns |
|  | Constant | -0.306 | 0.100 | 9.348 | 0.736 |  |  | 0.002** |
| UBR4-LRP1<br>was KD | OPHN1 KD | 0.044 | 0.119 | 0.139 | 1.045 | 0.828 | 1.321 | 0.709ns |
|  | Constant | -0.241 | 0.087 | 7.772 | 0.785 |  |  | 0.005** |

**Table 6.**
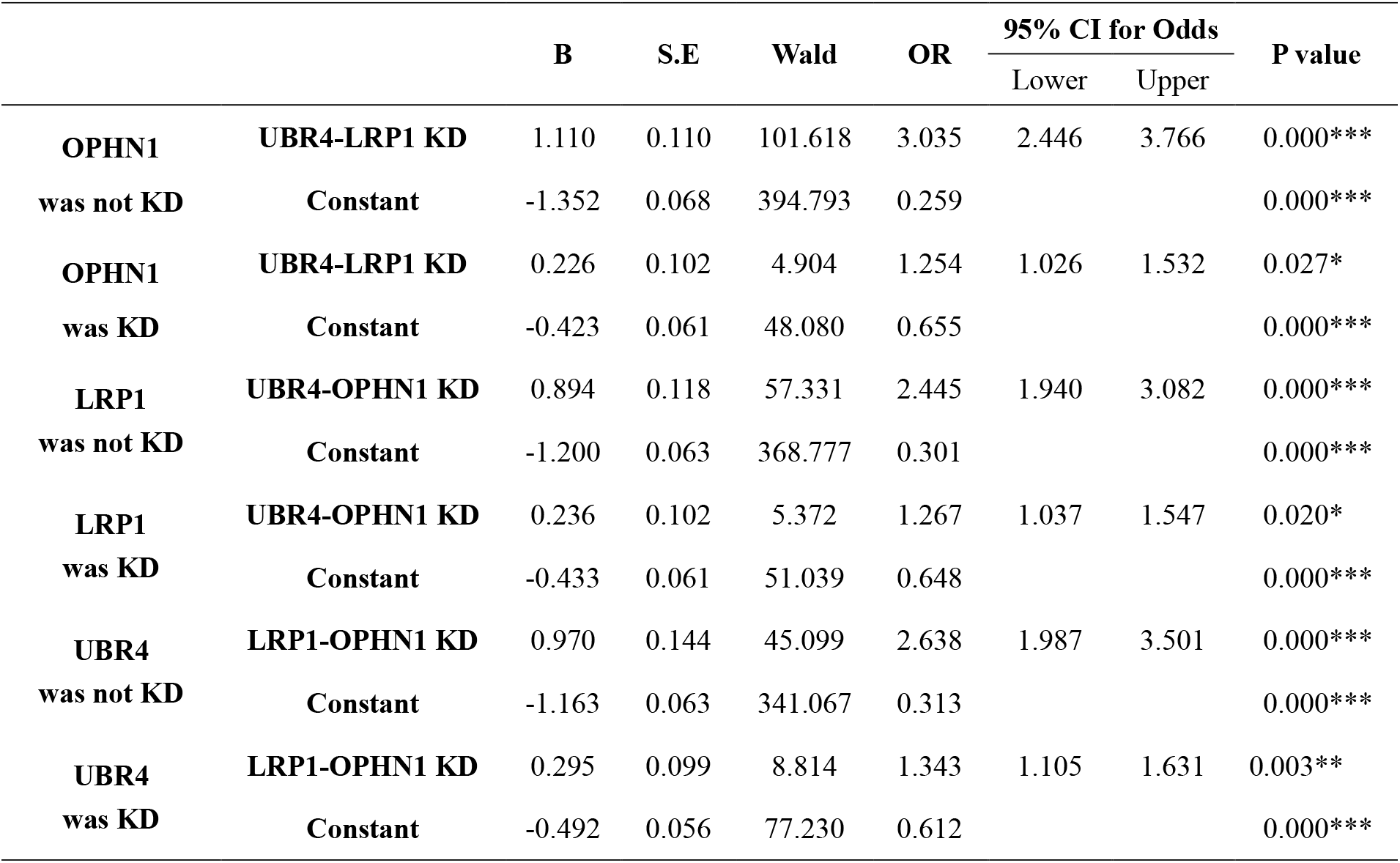
LR outcomes (seizures in relation to the digenic KDs)

#### 3.3. LRP1 KD served as main epileptogenic factor in the UBR4-LRP1 KD system

The UBR4-LRP1 double KD flies showed significantly higher seizure rate and longer recovery time than that of the single KD flies (Figure S1A). With a bit less significance, the LR equation for this digenic model was as follows:

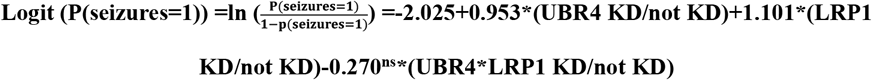

According to the LR result, UBR4 KD and LRP1 KD were both risk factors on seizures in the digenic KD system, while the negative interaction was insignificant even though the B value of the two genes had reduced when both genes were knockdown. Based on the two obviously larger B value of LRP1 KD, it was considered that LRP1 KD was the main epileptogenic factor in the UBR4-LRP1 digenic KD system. (Table S2-3)

#### 3.4. OPHN1 KD was main epileptogenic factor in the relevant digenic KD systems

The UBR4-OPHN1 and LRP1-OPHN1 double KD flies both showed significantly higher seizure rate than that of the single KD flies. The recovery time of OPHN1 KD flies was similar to the two relevant digenic KD flies while presented slower recover rate than the LRP1-OPHN1 KD flies. Besides, the seizure rate of OPHN1 KD flies was significantly higher than UBR4 KD flies in the UBR4-OPHN1 digenic KD system. (Figure S1B-C) The LR equation for this two digenic models were as follow:

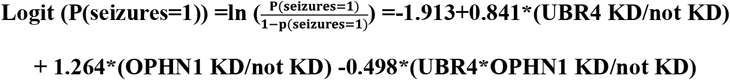

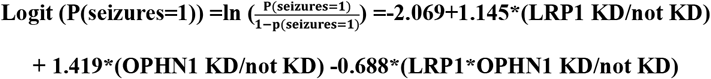

According to the LR result, UBR4 KD and OPHN1 KD were both risk factors on seizures in the UBR4-OPHN1 digenic KD system, so did the LRP1 KD and OPHN1 KD in the LRP1-OPHN1 digenic KD system. Moreover, negative interactions had been found significant in the two digenic KD systems that were relevant to the OPHN1 KD. Despite the existence of significant negative interactions, the monogenic KDs were still risk factors on seizures in the two digenic KD systems. OPHN1 KD was considered the main epileptogenic factor in both the UBR4-OPHN1 and LRP1-OPHN1 digenic KD system due to the four larger B value. (Table S5-6, S8-9)

## Discussion

Calcium ions are considered a significant factor in both epileptogenesis and seizure generation for the various ways in which calcium-dependent signaling pathways interact with neuronal activity(Steinlein, 2014). Besides, calcium overload is an important cause of neuronal apoptosis and secondary death in epilepsy(Gorji, 2001), and glutamate, as the main excitatory neurotransmitter in the central nervous system, can alter the permeability of the postsynaptic membrane to calcium ions, and abnormal inter-synaptic glutamate concentrations can cause a variety of neurological diseases, especially its persistently high level of seizures (Vyklicky et al., 2014). In our study, UBR4 KD, LRP1 KD, and OPHN1 KD flies showed significantly higher rate of epileptiform behaviors than the controls, indicating that the three candidate genes were epileptogenic. All these three genes have been discovered to be related to regulating calcium ion and glutamate concentrations.

UBR4, a gene encoding the ER- and microtubule-associated protein which are enriched in central nervous system neurons(Shim et al., 2008), plays a potential role in the vesicle trafficking of neuropeptides(Hegazi et al., 2022). Although there were no studies linking UBR4 to epilepsy, Camille Belzil et al discovered that the UBR4 deficiency leads the glutamate-induced Ca^2+^ influxes through the NMDA receptors, whose downstream carrying the aberrant calmodulin/calmodulin-dependent protein kinase II complex, then initiating a neuronal degenerative process of endoplasmic reticulum rupture and dysregulated calcium homeostasis(Belzil et al., 2013). Combined with our clinical data and through experimental validation, the epileptogenicity of UBR4 mutations can practically be verified now.

LRP1, a novel relevant gene of epilepsy, encodes the low-density lipoprotein receptor 1 which functions predominantly to mediate ligand endocytosis and modulates signal transduction processes(May et al., 2004). Epileptic seizures can be observed in LRP1-deficient flies and mice in previous experiments(Bres et al., 2020; Romeo et al., 2021; Zhang et al., 2022). According to the currently available research, LRP1 alters the expression of cellular glutamate transporter proteins and the LRP1 protein is very similar to the NMDA receptor in dendritic synapses, which may interact with the glutamatergic transmitter system to regulate the turnover and recirculation of synaptic proteins for synaptic regulation(May et al., 2004; Nakajima et al., 2013; Romeo et al., 2020). In addition, LRP1 is involved in the regulation of calcium homeostasis by affecting the expression of functional voltage-gated Ca^2+^ channels at the plasma membrane and increasing Ca^2+^ currents(Dolphin, 2012; Van Petegem et al., 2005). The epileptogenicity of LRP1 mutations was revalidated in our experiments again.

OPHN1(Oligophrenin 1, MIM:300127), is an identified human epilepsy-associated gene that encodes for a Rho-GTPase activating protein (RhoGAP). Some patients with mutations of OPHN1 were reported to exhibit seizures or epileptic syndromes(Bergmann et al., 2003; Zanni et al., 2005). OPHN1 protein could be involved in the regulation of the G protein cycle by down-regulating the RhoA/ Rho kinase signaling pathway to inhibit its inhibitory activity on synaptic vesicle cycling and AMPAR internalization(Khelfaoui et al., 2007). Moreover, due to the proline-rich structural domains and endocytophilic proteins, OPHN1 may also regulate endocytosis by coordinating actin and membrane dynamics(Nadif Kasri et al., 2011; Nakano-Kobayashi et al., 2009), which in turn work together to achieve regulation of the glutamatergic vesicle endocytosis cycle and excitatory synaptic transmission. The epileptogenicity of the OPHN1 mutation was validated at the whole animal level by our experiments.

It was estimated that approximately 70-80% of epilepsy occurrences are estimated to be triggered by one or more genetic factors(Myers & Mefford, 2015). An experiment conducted on mice demonstrated that compared to single mutants, double heterozygotes exhibited significantly more severe symptoms in terms of age of onset, types, and severity of epilepsy(Kearney et al., 2006). When we further explored the polygenic effects, it also showed that the seizure rates of the three groups of digenic KD flies and the trigenic KD flies were significantly higher than that of the corresponding monogenic KD flies. However, flies of the trigenic KD did not show stronger seizures than those of the digenic KD, implying the possible existence of some negative interaction effects.

According to the results of the logistic regression analysis that we used to investigate the interactions and effects, monogenic KDs of the three genes were risk factors for seizures in the trigenic KD system, meanwhile, significant negative interactions were found in the UBR4*OPHN1 KD and LRP1*OPHN1 KD. Thus, it explains why the seizures in the trigenic KD model are similar to those in the digenic KD model. After analysis in more details, the effects of the three monogenic KDs were insignificant while the promotive effects of the three digenic KDs when the three genes were knockdown were significant despite the negative interactions. Epileptic behavior was considered to be triggered mainly by digenic KDs together with their interaction when UBR4-LRP1-OPHN1 was KD. Taking all the results of logistic regression analysis together, LRP1-OPHN1 KD was regarded as the main causative genetic factor for seizures due to the relatively higher B values in the trigenic LR model. Moreover, UBR4 KD was hypothesized to play a protective role against seizures in the trigenic KD system, nevertheless, related mechanistic studies need to be done to further substantiate this conclusion.

It is speculated that in the trigenic knockdown Drosophila system we constructed, negative interactions could be generated mainly by the co-regulation of inter-synaptic glutamate concentration which could indirectly affect calcium homeostasis. Moreover, the possible protective effect against seizures of UBR4 KD might be based on the direct co-regulation of Ca^2+^ homeostasis by UBR4 KD and LRP1 KD which compensates for the dysregulation of calcium homeostasis that promotes epileptogenesis. Therefore, calcium imaging experiments in the brain tissue of Drosophila will be conducted next to preliminarily explore the individual and interaction mechanisms among these three genes, and then further find possible therapeutic targets for the disease.

## Author Contributions

Jing-Da Qiao designed the study. Rui-Na Huang, Si-Yuan Luo, Tao-Huang, Xiong-Sheng Li, Fan-Chao Zhou, Ling-Ying Li and Jing-Da Qiao performed the experiments. Rui-Na Huang, Xiong-Sheng Li, Fan-Chao Zhou and Ze-Ru Chen analyzed the data. Rui-Na Huang, Si-Yuan Luo, Tao-Huang and Jing-Da Qiao wrote the manuscript.

## Acknowledgments

We thank Tsinghua Fly Center for donating transgenic RNAi lines. This study was funded by Guangzhou Medical University (Funding No. 02-408-240603131147 to Rui-Na Huang), the Science and Technology Project of Guangzhou (Grant No. 2024A04J3396 to Jing-Da Qiao), the Guangdong Basic and Applied Basic Research Foundation (Grant No. 2022A1515111123 to Jing-Da Qiao), and the National Natural Science Foundation of China (Grant No. 82071548 to Bin Tao)

