## Supplemental Data for "The Interaction of *UBR4, LRP1*, and *OPHN1* in Refractory Epilepsy: *Drosophila* Model to Investigate the Oligogenic Effect on Epilepsy"

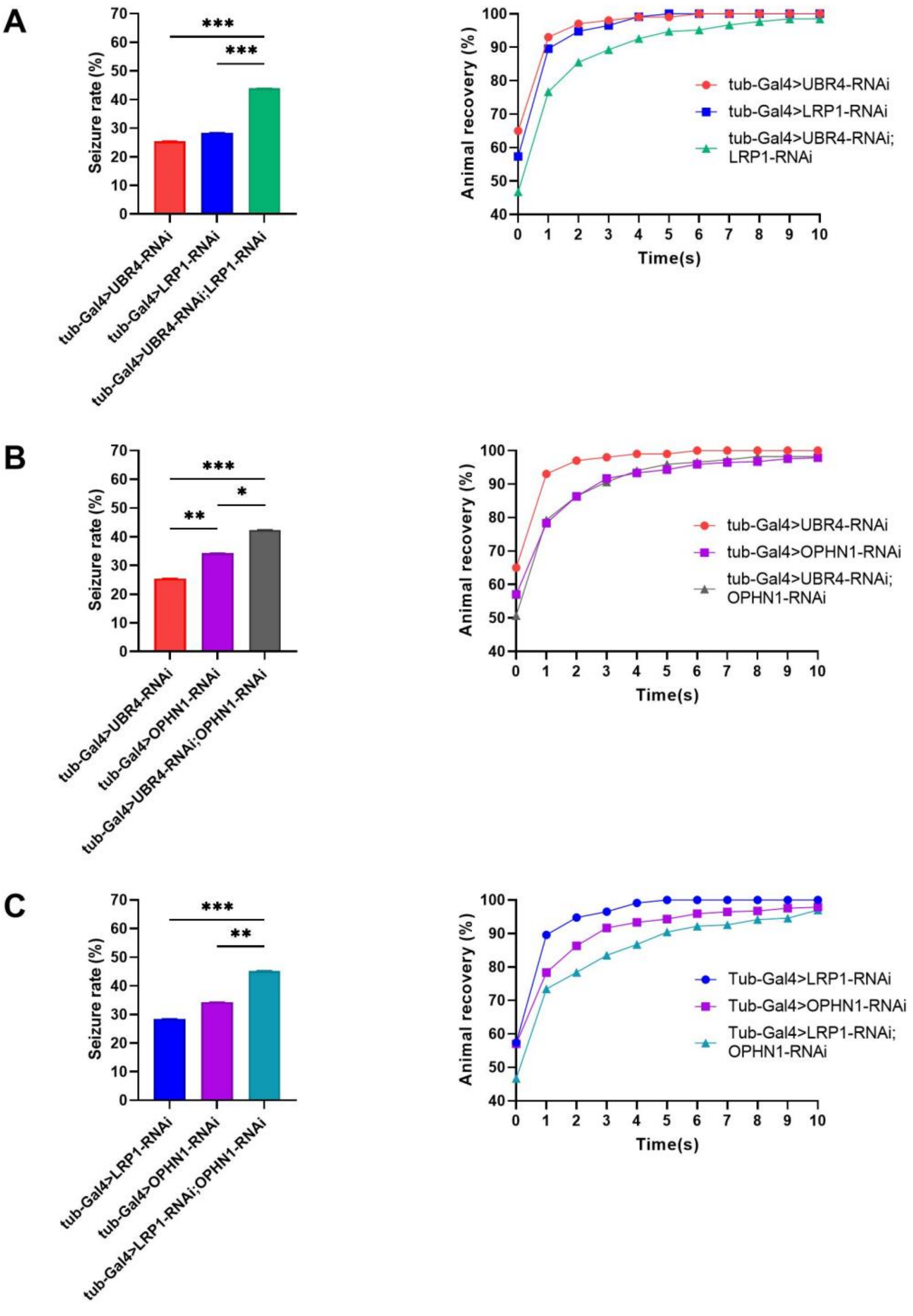


Figure S1. Seizure behavior of digenic KD flies compared with monogenic KD. (A) Seizure rates and recovery time of *UBR4* KD, *LRP1* KD, and *UBR4-LRP1* KD flies. (B) Seizure rates and recovery time of *UBR4* KD, *OPHN1* KD, and *UBR4-OPHN1* KD flies. (C) Seizure rates and recovery time of *LRP1* KD, *OPHN1* KD, and *LRP1-OPHN1* KD flies. Error bars represent 95% Confidence Interval (CI) of seizure rates.

Table S1. Total seizures and the KD of two genes (*UBR4*&*LRP1*)

| Group | seizures | UBR4 KD | LRP1 KD | N |
|---|---|---|---|---|
| **Double KD** | 1 | 1 | 1 | 238 |
| | 0 | 1 | 1 | 303 |
| **UBR4 KD** | 1 | 1 | 0 | 104 |
| | 0 | 1 | 0 | 304 |
| **LRP1 KD** | 1 | 0 | 1 | 102 |
| | 0 | 0 | 1 | 257 |
| **Control** | 1 | 0 | 0 | 57 |
| | 0 | 0 | 0 | 432 |

Table S2. LR outcomes (interaction between *UBR4* and *LRP1*)

| | B | S.E | Wald | OR | 95% CI for Odds | | P value |
|---|---|---|---|---|---|---|---|
| | | | | | Lower | Upper | |
| **UBR4 KD** | 0.953 | 0.181 | 27.705 | 2.593 | 1.818 | 3.697 | 0.000*** |
| **LRP1 KD** | 1.101 | 0.183 | 36.145 | 3.008 | 2.101 | 4.307 | 0.000*** |
| **UBR4*LRP1** | -0.270 | 0.232 | 1.352 | 0.763 | 0.484 | 1.203 | 0.245ns |
| **Constant** | -2.025 | 0.141 | 206.567 | 0.132 | | | 0.000*** |

Table S3. LR outcomes (seizures in relation to *UBR4* KD or *LRP1* KD)

| | | B | S.E | Wald | OR | 95% CI for Odds | | P value |
|---|---|---|---|---|---|---|---|---|
| | | | | | | Lower | Upper | |
| **LRP1 was not KD** | **UBR4 KD** | 0.953 | 0.181 | 27.705 | 2.593 | 1.818 | 3.697 | 0.000*** |
| | **Constant** | -2.025 | 0.141 | 206.567 | 0.132 | | | 0.000*** |
| **LRP1 was KD** | **UBR4 KD** | 0.683 | 0.146 | 21.984 | 1.979 | 1.488 | 2.633 | 0.000*** |
| | **Constant** | -0.924 | 0.117 | 62.356 | 0.397 | | | 0.000*** |
| **UBR4 was not KD** | **LRP1 KD** | 1.101 | 0.183 | 36.145 | 3.008 | 2.101 | 4.307 | 0.000*** |
| | **Constant** | -2.025 | 0.141 | 206.567 | 0.132 | | | 0.000*** |
| **UBR4 was KD** | **LRP1 KD** | 0.831 | 0.143 | 33.854 | 2.296 | 1.735 | 3.038 | 0.000*** |
| | **Constant** | -1.073 | 0.114 | 89.156 | 0.342 | | | 0.000*** |

Table S4. Total seizures and the KD of two genes (*UBR4&OPHN1*)

| Group | seizures | UBR4 KD | OPHN1 KD | N |
|---|---|---|---|---|
| **Double KD** | 1 | 1 | 1 | 173 |
| | 0 | 1 | 1 | 235 |
| **UBR4 KD** | 1 | 1 | 0 | 104 |
| | 0 | 1 | 0 | 304 |
| **OPHN1 KD** | 1 | 0 | 1 | 163 |
| | 0 | 0 | 1 | 312 |
| **Control** | 1 | 0 | 0 | 40 |
| | 0 | 0 | 0 | 271 |

Table S5. LR outcomes (interaction between *UBR4* and *OPHN1*)

| | B | S.E | Wald | OR | 95% CI for Odds | | P value |
|---|---|---|---|---|---|---|---|
| | | | | | Lower | Upper | |
| **UBR4 KD** | 0.841 | 0.204 | 16.988 | 2.318 | 1.554 | 3.457 | 0.000*** |
| **OPHN1 KD** | 1.264 | 0.195 | 42.010 | 3.540 | 2.415 | 5.187 | 0.000*** |
| **UBR4*OPHN1** | -0.498 | 0.247 | 4.062 | 0.608 | 0.375 | 0.986 | 0.044* |
| **Constant** | -1.913 | 0.169 | 127.587 | 0.148 | | | 0.000*** |

Table S6. LR outcomes (seizures in relation to *UBR4* KD or *OPHN1* KD)

| | | B | S.E | Wald | OR | 95% CI for Odds | | P value |
|---|---|---|---|---|---|---|---|---|
| | | | | | | Lower | Upper | |
| **OPHN1 was not KD** | **UBR4 KD** | 0.841 | 0.204 | 16.988 | 2.318 | 1.554 | 3.457 | 0.000*** |
| | **Constant** | -1.913 | 0.169 | 127.587 | 0.148 | | | 0.000*** |
| **OPHN1 was KD** | **UBR4 KD** | 0.343 | 0.139 | 6.071 | 1.409 | 1.073 | 1.851 | 0.014* |
| | **Constant** | -0.649 | 0.097 | 45.131 | 0.522 | | | 0.000*** |
| **UBR4 was not KD** | **OPHN1 KD** | 1.264 | 0.195 | 42.010 | 3.540 | 2.415 | 5.187 | 0.000*** |
| | **Constant** | -1.913 | 0.169 | 127.587 | 0.148 | | | 0.000*** |
| **UBR4 was KD** | **OPHN1 KD** | 0.766 | 0.151 | 25.600 | 2.152 | 1.599 | 2.896 | 0.000*** |
| | **Constant** | -1.073 | 0.114 | 89.156 | 0.342 | | | 0.000*** |

Table S7. Total seizures and the KD of two genes (*LRP1&OPHN1*)

| Group | seizures | LRP1 KD | OPHN1 KD | N |
|---|---|---|---|---|
| **Double KD** | 1 | 1 | 1 | 108 |
| | 0 | 1 | 1 | 131 |
| **LRP1 KD** | 1 | 1 | 0 | 102 |
| | 0 | 1 | 0 | 257 |
| **OPHN1 KD** | 1 | 0 | 1 | 163 |
| | 0 | 0 | 1 | 312 |
| **Control** | 1 | 0 | 0 | 35 |
| | 0 | 0 | 0 | 277 |

Table S8. LR outcomes (interaction between *LRP1* and *OPHN1*)

| | B | S.E | Wald | OR | 95% CI for Odds | | P value |
|---|---|---|---|---|---|---|---|
| | | | | | Lower | Upper | |
| **LRP1 KD** | 1.145 | 0.214 | 28.556 | 3.141 | 2.064 | 4.780 | 0.000*** |
| **OPHN1 KD** | 1.419 | 0.204 | 48.523 | 4.135 | 2.773 | 6.164 | 0.000*** |
| **LRP1*OPHN1** | -0.688 | 0.269 | 6.571 | 0.502 | 0.297 | 0.850 | 0.010* |
| **Constant** | -2.069 | 0.179 | 132.977 | 0.126 | | | 0.000*** |

Table S9. LR outcomes (seizures in relation to *LRP1* KD or *OPHN1* KD)

| | | B | S.E | Wald | OR | 95% CI for Odds | | P value |
|---|---|---|---|---|---|---|---|---|
| | | | | | | Lower | Upper | |
| **OPHN1 was not KD** | **LRP1 KD** | 1.145 | 0.214 | 28.556 | 3.141 | 2.064 | 4.780 | 0.000*** |
| | **Constant** | -2.069 | 0.179 | 132.977 | 0.126 | | | 0.000*** |
| **OPHN1 was KD** | **LRP1 KD** | 0.456 | 0.162 | 7.933 | 1.578 | 1.149 | 2.168 | 0.005** |
| | **Constant** | -0.649 | 0.097 | 45.131 | 0.522 | | | 0.000*** |
| **LRP1 was not KD** | **OPHN1 KD** | 1.419 | 0.204 | 48.523 | 4.135 | 2.773 | 6.164 | 0.000*** |
| | **Constant** | -2.069 | 0.179 | 132.977 | 0.126 | | | 0.000*** |
| **LRP1 was KD** | **OPHN1 KD** | 0.731 | 0.175 | 17.472 | 2.077 | 1.474 | 2.927 | 0.000*** |
| | **Constant** | -0.924 | 0.117 | 62.356 | 0.397 | | | 0.000*** |
